# Fast volumetric mapping of human brain slices

**DOI:** 10.1101/2020.10.27.357186

**Authors:** Luca Pesce, Annunziatina Laurino, Vladislav Gavryusev, Giacomo Mazzamuto, Giuseppe Sancataldo, Marina Scardigli, Matteo Roffilli, Ludovico Silvestri, Irene Costantini, Francesco Saverio Pavone

**Affiliations:** Department of Physics and Astronomy, University of Florence Via G. Sansone, 1 – 50019 Sesto Fiorentino (FI), Italy; European Laboratory for Non-linear Spectroscopy (LENS), University of Florence Via Nello Carrara, 1 – 50019 Sesto Fiorentino (FI), Italy; National Institute of Optics, National Research Council; Bioretics Srl, Cesena, Italy

**Keywords:** Light sheet microscopy, whole brain imaging, quantitative imaging, tissue clearing, human brain

## Abstract

We still lack a detailed map of the anatomical disposition of neurons in the human brain. A complete map would be an important step for deeply understanding the brain function, providing anatomical information useful to decipher the neuronal pattern in healthy and diseased conditions. Here, we present several important advances towards this goal, obtained by combining a new clearing method, advanced Light Sheet Microscopy and automated machinelearning based image analysis. We perform volumetric imaging of large sequentially stained human brain slices, labelled for two different neuronal markers NeuN and GAD67, discriminating the inhibitory population and reconstructing the brain connectivity.

## 1. Introduction

One of the most important challenges in today’s neuroscience is the volumetric mapping of the neuronal architecture in human brain samples. Recent advances in tissue clearing techniques have enabled to access the structure and function of large intact biological tissues [1, 2]. These approaches use several chemicals capable of dissolving cellular membrane lipids to render the tissue completely transparent by homogenizing the refractive index. Different optical clearing agents have been developed in order to reduce the scattering of the biological specimen, enhance the contrast image and increase imaging penetration depth [3, 4]. Furthermore, new methods capable of embedding and expanding the sample in an acrylamide hydrogel allow enhancing the final optical resolution, at the cost of an increased sample size (4 – 20 times expansion) [5]. In addition, on the imaging side, Light Sheet Fluorescence Microscopy provides with high resolution the multiscale information required to investigate the characteristics of neuronal tissues [6, 7]. Today, by combining these two optical approaches, scientists are able to accelerate discoveries in life sciences and reconstruct the mammalian brain architecture. However, obtaining micrometrically resolved anatomical information over large portions of post-mortem human brain is still a challenge in biology. Indeed, the structural properties of the tissue, the variability of the sample’s post-mortem condition and the impossibility to use genetically encoded fluorescent probes, make the treatment of such a sample extremely difficult and laborious. Here, we present the volumetric imaging of sequentially stained human brain slices, labelled for NeuN and GAD67 and acquired using a custom dual-view inverted confocal light sheet fluorescence microscope (di2CLSFM). Our instrument, by fusing the two orthogonal views, enables an isotropic resolution of 1 μm along the three optical axes (x, y and z). By preserving the molecular disposition of the biomolecules through the SWITCH method [8] and by clarifying with the 2’2-Thiodiethanol agent [9], we are able to discriminate the inhibitory population in the human cortex and to reconstruct the brain connectivity over large sample volumes, leveraging automated machinelearning based image analysis. Also, we show the complete pipeline developed to investigate the cytoarchitecture characteristics of the human cortex, from the sample treatment to the quantitative analysis.

## 2. Methodology

As shown in Fig. 2, we develop a specific pipeline in order to analyze large volume of human brain sample. Large portion of the samples are cut into slices with 500 μm thick. To clarify the brain slices, we use an adapted protocol from Murray et al [6] called SWITCH. This method allows diffusing and crosslinking glutaraldehyde (GA) in the sample, by changing the pH value in order to stabilize the endogenous proteins. Then, the transformed tissue is treated with strong detergent [sodium dodecyl sulfate (SDS)] for lipid removal. In this way, we avoid loss of the protein content in the specimen, necessary for the antibody recognition. Briefly, the specimen is incubated in a Switch off solution, consisting of 50% PBS titrated to pH 3 using HCl, 25% 0.1 M HCl, 25% 0.1 M potassium hydrogen phthalate (KHP) and 4% glutaraldehyde. After 24 h, the solution is replaced with PBS pH 7.6 with 1% glutaraldehyde. Finally, the samples are incubated in the clearing solution for 7 d at 53 °C. After washing with PBS + 0.1% Triton (PBST) for 24 h, the primary antibodies are incubated at 4 °C for 24 h in PBST (dilution for NeuN and Gad67 are 1:50 and 1:200, respectively). Following several washes in PBST, the samples are incubated with the secondary antibodies (dilution 1:200). After fixation with 4% PFA, the slices are rendered transparent by soaking the samples in increasing solution of 20%, 40% and 68% (vol/vol) of 2,2’-thiodiethanol in PBS, and imaged with our custom di2CLSFM. This step permits to achieve high penetration depth and avoid any optical aberration by matching the refractive index of our biological samples.

**Figure 1.**
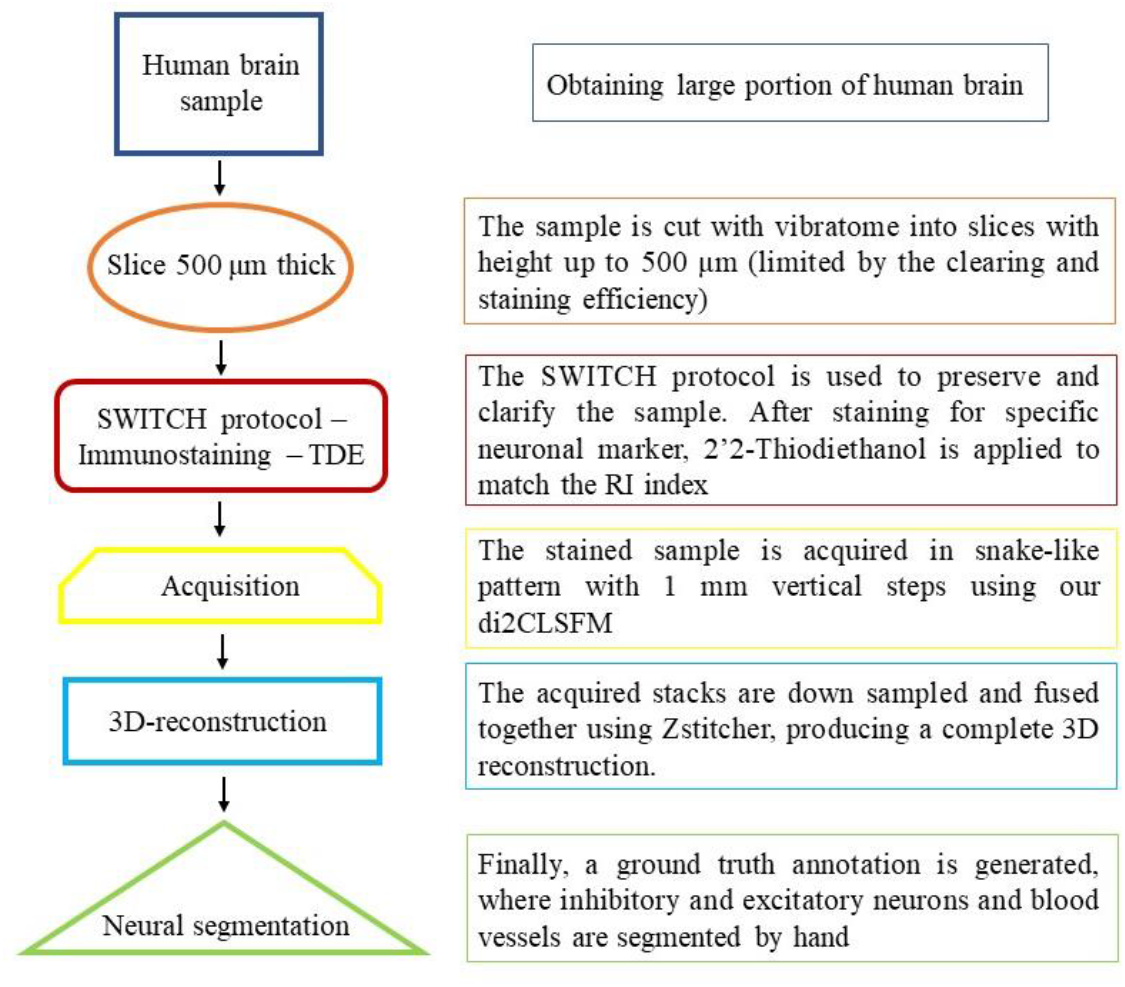
Schematic representation of our pipeline developed to analyze large portion of the human brain. The 500 μm thick cortical slices are clarified, labelled and acquired using di2CLSFM. Next, to evaluate the cytoarchitecture of our sample, a fast down sampled 3D reconstruction is performed using Zsticher. Finally, we are developing a ground truth annotation using the high-resolution images, by generating three different classes: excitatory and inhibitory neurons and vessels.

**Figure 2.**
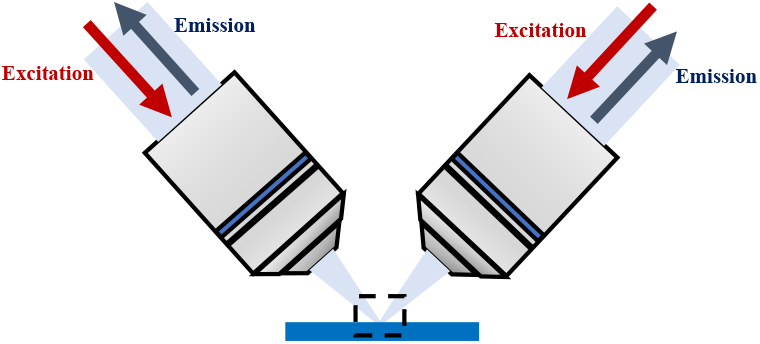
Schematic illustration of the di2CLSFM dual-view inverted arrangement, that allows sequential mosaic imaging of extended samples with size up to 300×300×0.8 mm^3^. Multiple labelled human brain samples can be held in one or more custom slide holders, fully immersed in the refractive index matching immersion solution.

Fig. 2 shows a schematic illustration of our custom-made light sheet microscope (di2CLSFM). Both objectives alternately excite and collect the fluorescence signal. Their characteristics are 12x magnification, NA 0.53, WD 10 mm, with a correction collar for refractive index matching with the immersion solution. Additionally, the set-up includes 4 overlapped laser lines (405, 488, 561 and 633 nm) with the corresponding fluorescence filter bands, to detect up to four differently stained structures. A 50/50% beam splitter conveys the combined beam into the two identical optical excitation pathways, where an acousto-optical-tunable-filter modulates the transmitted power and wavelength and a galvo mirror sweeps it across the detection focal plane, generating a digitally scanned light sheet. The induced fluorescence is collected by each objective onto a Hamamatsu ORCA Flash4.0v3 sCMOS camera. The sample is imaged by translating it along the horizontal direction in a snake-like pattern.

## 3. Data

A representative example of high-speed quantitative imaging of large intact biological specimens labelled for NeuN is presented in Fig. 3. Furthermore, by optimizing the image contrast, we demonstrate the possibility to perform a 2-colour immunolabeling. Fig. 4 shows the high-resolution imaging of human brain slices labelled for NeuN (A) and GAD67 (B) with Alexa Fluor 488 and Alexa Fluor 647, respectively. In this way, it is possible to reconstruct the neuronal pattern in large human samples (Fig. 3, A) and discriminate the inhibitory neurons (Fig. 4, B).

**Figure 3.**
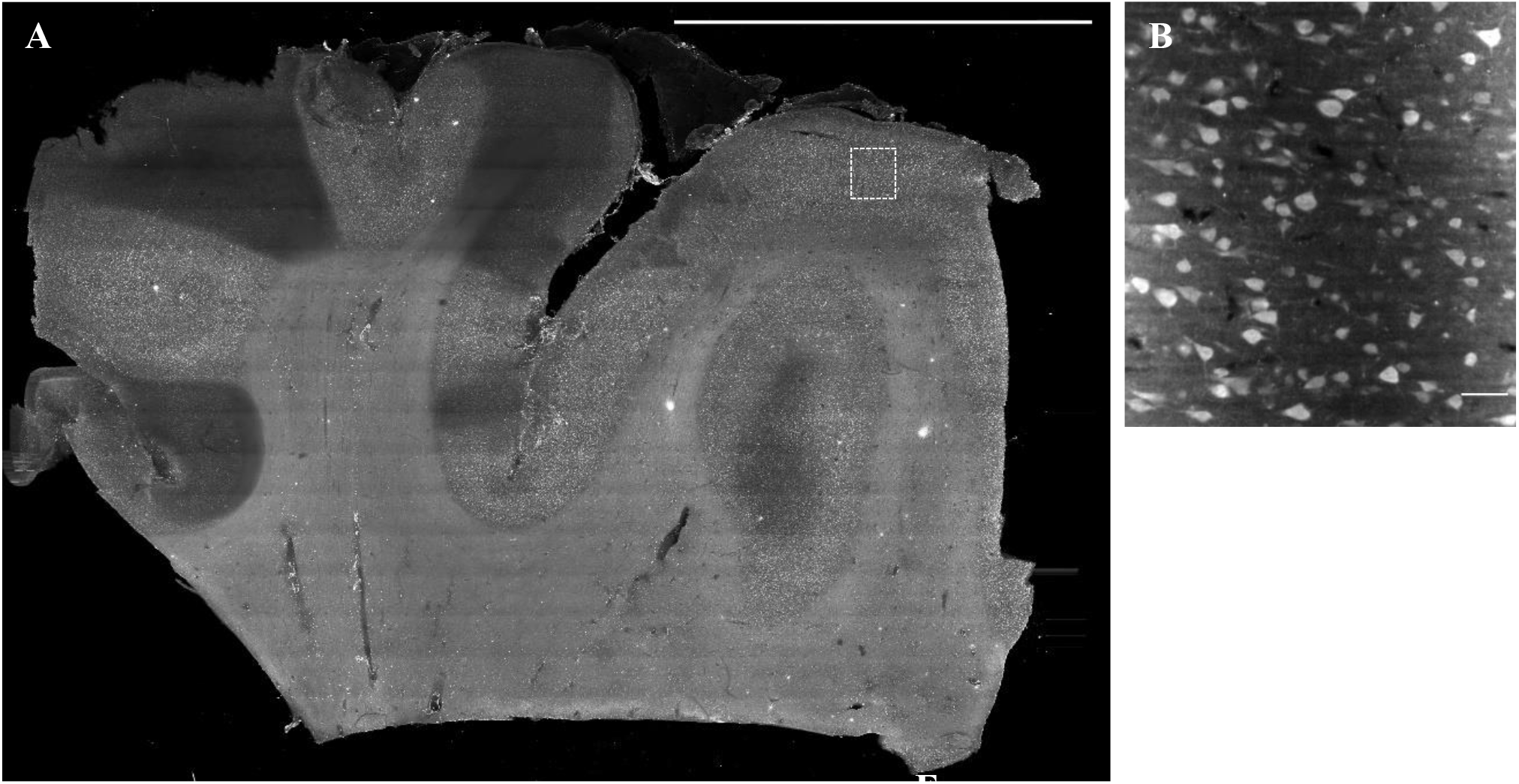
The images A and B show, respectively, a downscaled reconstruction and the corresponding high-resolution image of the neuronal localization in a slice of a large human brain portion labelled for NeuN with Alexa 647. Scale bar 1 cm and 100 μm, respectively.

**Figure 4.**
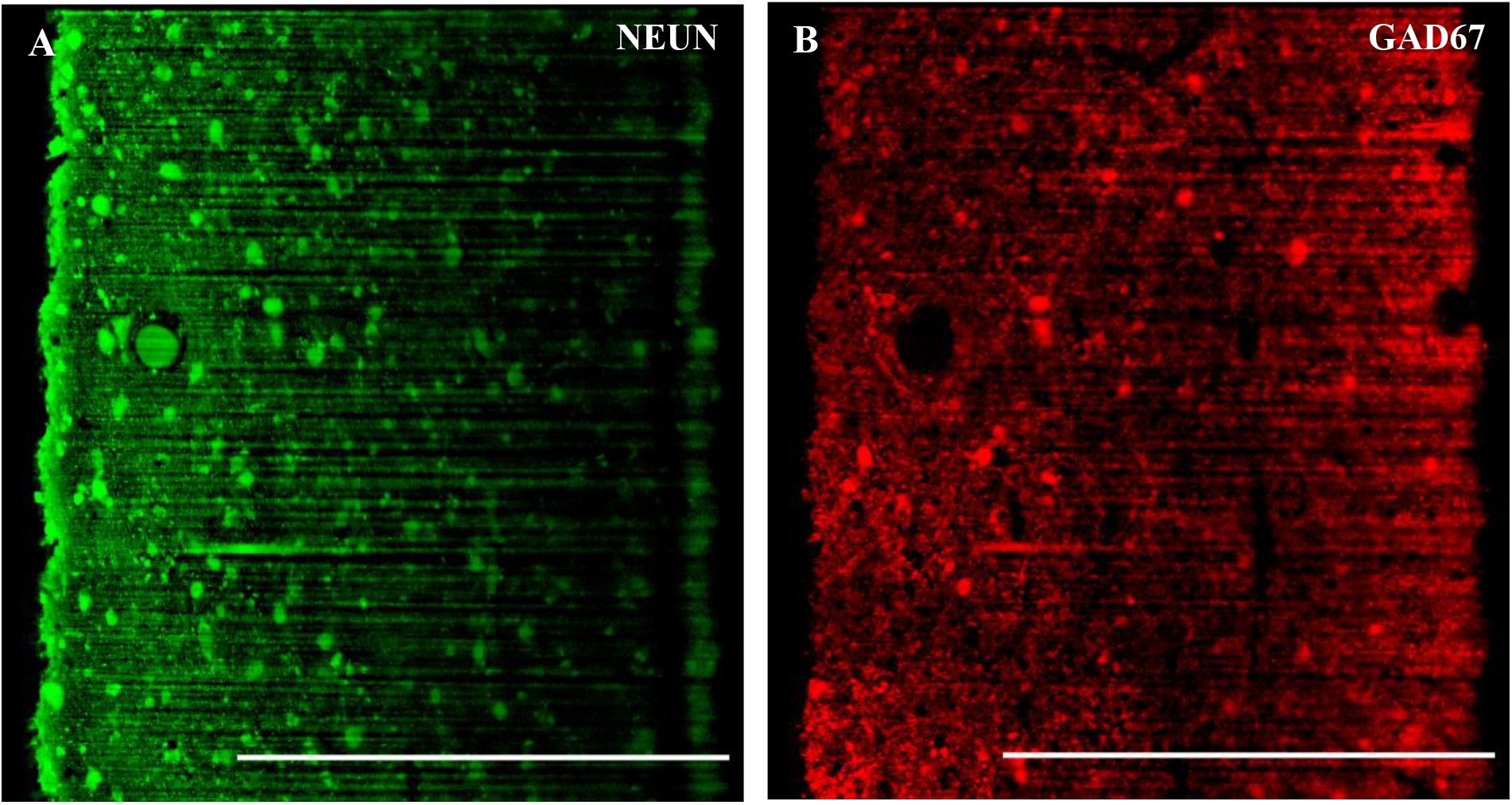
Costaining with NeuN (A), which labels all neurons, and Gad67 (B), which labels only the inhibitory population. Scale bar 500 μm.

## 4. Conclusion

The high-contrast provided by our optimized immunolabeling protocol, the deepened light penetration (Fig 4 and 5) and the di2CLSFM confocal detection improve the performance of our automatic segmentation algorithm, based on machinelearning and capable of handling datasets exceeding one TeraByte. Such strategies, already exploited in neuronal segmentation [4, 11], are well suited for revealing the organization of neuronal tissue both in healthy and diseased condition.

## 5. Acknowledgments

This project has received funding from the European Union’s Horizon 2020 Framework Programme for Research and Innovation under Grant Agreement No. 720270 (Human Brain Project SGA1), No. 785907 (Human Brain Project SGA2) and the NIH BRAIN Initiative and Marie Skłodowska-Curie Grant Agreement No. 793849 (MesoBrainMicr).

